# Mechanical coordination between anaphase A and B drives asymmetric chromosome segregation

**DOI:** 10.1101/2025.10.02.680008

**Authors:** Ana M. Dias Maia Henriques, Tim Davies, Serge Dmitrieff, Nicolas Minc, Julie C. Canman, Julien Dumont, Gilliane Maton

## Abstract

Chromosome segregation during anaphase occurs through two mechanistically distinct processes: anaphase A, in which chromosomes move toward spindle poles, and anaphase B, in which the anaphase spindle elongates through cortical astral microtubule pulling forces. *Caenorhabditis elegans* embryos have been thought to rely primarily on anaphase B, with little to no contribution from anaphase A. Here, we uncover a novel anaphase A mechanism in *C. elegans* embryos, driven by the kinesin-13 KLP-7^MCAK^ and opposed by the kinesin-12 KLP-18. We found that the extent of chromosome segregation during anaphase A is asymmetrically regulated by cell polarity cues and modulated by mechanical tension within the spindle, generated by opposing forces acting on chromosomes and spindle poles. Additionally, we found that the contribution of anaphase A to chromosome segregation increases progressively across early embryonic divisions. These findings uncover an unexpected role for anaphase A in early *C. elegans* development and reveal a KLP-7^MCAK^-dependent mechanical coordination between anaphase A and anaphase B driven chromosome segregation.

**eTOC summary:** Dias Maia Henriques et al. uncover an anaphase A pathway, driven by the kinesin-13 KLP-7 and opposed by the kinesin-12 KLP-18, that contributes to chromosome segregation in early *C. elegans* embryos. Its activity is regulated by spindle tension, cell polarity cues, and progressively increases during early embryonic divisions.

## INTRODUCTION

During cell division, faithful segregation of chromosomes between daughter cells is crucial for maintaining genome integrity. Accurate chromosome segregation depends on kinetochore-mediated attachments to dynamic spindle microtubules. Multiprotein kinetochores translate the dynamic behavior of microtubules into coordinated chromosome movements, ensuring their alignment at the spindle equator during metaphase and their subsequent physical separation into two equal sets of sister chromatids during anaphase (Ariyoshi and Fukagawa, 2023; Musacchio and Desai, 2017).

After cohesin cleavage, sister chromatid separation during anaphase is governed by two distinct processes, anaphase A and anaphase B, which can occur independently, simultaneously, in any sequence, and to varying extents, depending on cell types and organisms (McIntosh, 2021; Vukusic and Tolic, 2021). Anaphase A is characterized by chromosome movements toward the spindle poles through the shortening of kinetochore microtubules (Asbury, 2017). Anaphase B involves the separation of spindle poles, resulting in spindle elongation (Scholey et al., 2016).

During anaphase B, spindle elongation can be driven by spindle-external forces exerted through astral microtubules pulling on spindle poles from the cell cortex, or by spindle-internal pushing forces emanating from the spindle midzone or central spindle microtubules (McIntosh et al., 2012; Roostalu et al., 2010). In the cortical pulling mechanism, considered as the primary force-driving mechanism in *C. elegans* embryos, astral microtubules anchored at both the cortex and spindle poles transmit forces to the poles, pulling the associated kinetochore microtubules and attached chromatids apart (Labbe et al., 2004; Oegema et al., 2001). Anaphase B cortical pulling forces rely on minus-end-directed microtubule motors, such as dynein anchored at the cell cortex, or on the depolymerization of cortically anchored astral microtubule plus-ends (Grill et al., 2001; Grishchuk et al., 2005; Guild et al., 2017). The pushing mechanism of anaphase B relies on midzone and/or interpolar antiparallel microtubules that push directly onto the spindle poles or indirectly on microtubules emanating from the spindle poles (Vukusic et al., 2019; Ward et al., 2014). Pushing can be exerted through the antiparallel sliding or plus-end polymerization of interpolar midzone or bridging fiber microtubules (Brust-Mascher et al., 2009; Vukusic et al., 2017).

During anaphase A, the sites of kinetochore microtubule depolymerization and tubulin subunit loss vary significantly between cell types and can even change dynamically within a single cell over time. In most fungi, for instance, kinetochore microtubule shortening occurs primarily at kinetochores, where microtubule depolymerization drives chromosome movement via a ’Pac-Man’ mechanism (Gorbsky et al., 1987; Maddox et al., 2000; Mallavarapu et al., 1999). In contrast, in many animal and plant cells, kinetochore microtubules exhibit continuous poleward flux during metaphase, characterized by simultaneous polymerization at kinetochores and depolymerization at spindle poles throughout metaphase (Dhonukshe et al., 2006; LaFountain et al., 2001; Mitchison and Salmon, 2001). At anaphase onset, microtubule polymerization at kinetochores slows or ceases. Thus anaphase A chromosome movement is, in part, a continuation of metaphase microtubule flux, without the counterbalance of kinetochore-associated microtubule polymerization (Mallavarapu et al., 1999). Both mechanisms can operate simultaneously, as observed in mitotic human cells, where chromosome movement during anaphase A results from a combination of ’Pac-Man’ and flux-based kinetochore microtubule shortening (Ganem et al., 2005; Zhang et al., 2007). The depolymerization of kinetochore microtubules can be driven by microtubule-depolymerizing kinesin-8 or kinesin-13 family members, or by microtubule-severing enzymes of the spastin, fidgetin or katanin families acting either at spindle poles or directly at kinetochores (Gupta et al., 2006; Maney et al., 1998; Rogers et al., 2004; Walczak et al., 1996; Zhang et al., 2007). Additionally, recent studies have revealed that kinetochore microtubules can integrate into the spindle without extending all the way to the centrosomes (Kiewisz et al., 2022; Yu et al., 2019). These short microtubules could also contribute to anaphase A motion of chromosomes through poleward motor-driven parallel sliding along non-kinetochore microtubules (Hueschen et al., 2017; Sikirzhytski et al., 2014).

Despite progress in identifying key molecular players and mechanisms, the precise regulation and interplay of microtubule depolymerization, flux, and sliding in anaphase A are still not fully understood. Furthermore, the coordination between anaphase A and anaphase B forces remains largely unexplored. While both anaphase A and B contribute to chromosome segregation, how they are temporally and mechanistically integrated to regulate chromosome separation is unclear. By analyzing chromosome movements during anaphase in *C. elegans* embryos, we found that, although chromosome segregation is primarily driven by an anaphase B mechanism, anaphase A contributes significantly to the overall separation of sister chromatids. Our results show that the cell polarity machinery asymmetrically regulates anaphase A on the two halves of the spindle in the asymmetrically dividing zygote. We further demonstrate that anaphase A chromosome movement is driven by the kinesin-13 KLP-7^MCAK^ and regulated by spindle tension, which arises from astral microtubule cortical pulling forces, driving anaphase B, opposed by the anaphase central spindle. Finally, analysis of multicellular early embryos revealed that anaphase A increasingly contributes to overall sister chromatid separation across successive cleavage divisions during early development. Our study unveils unexpected insights into the coordination between anaphase A and B and their mechanistic integration during chromosome segregation.

## RESULTS/DISCUSSION

### Chromosomes undergo biphasic movement within the spindle during anaphase in the C. elegans zygote

To characterize chromosome movements during anaphase in the *C. elegans* zygote, we developed a semi-automated, high-resolution, 4D-tracking assay for poles and chromosomes during the first embryonic division. We performed confocal microscopy on live embryos co-expressing a chromosome (mCherry::HIS-58^H2B^) and spindle pole (GFP::TBG-1^γ-tubulin^) marker, as well as a GFP-tagged AIR-2^AuroraB^ kinase, as a marker of anaphase onset and central spindle integrity (**Fig. 1 A** and **Video 1**) (Dumont and Maton, 2025; Maton et al., 2015). We measured the distance between the spindle poles (pole separation), and between the two sets of segregating chromosomes (chromosome segregation) over time (**Fig. 1 B**) (Edwards et al., 2018). Surprisingly, the two profiles were not strictly parallel, as one might expect if anaphase B was the sole driver of chromosome segregation in *C. elegans* zygotes (Labbe et al., 2004) (**Fig. 1 C**). Instead, we found that chromosomes segregated significantly faster than the spindle poles separated during the first 100 seconds following anaphase onset. This observation highlights the existence of a chromosome movement within the spindle—referred to hereafter as chromosome displacement—that occurs independently of spindle pole separation (**Fig. 1 B**).

**Figure 1.**
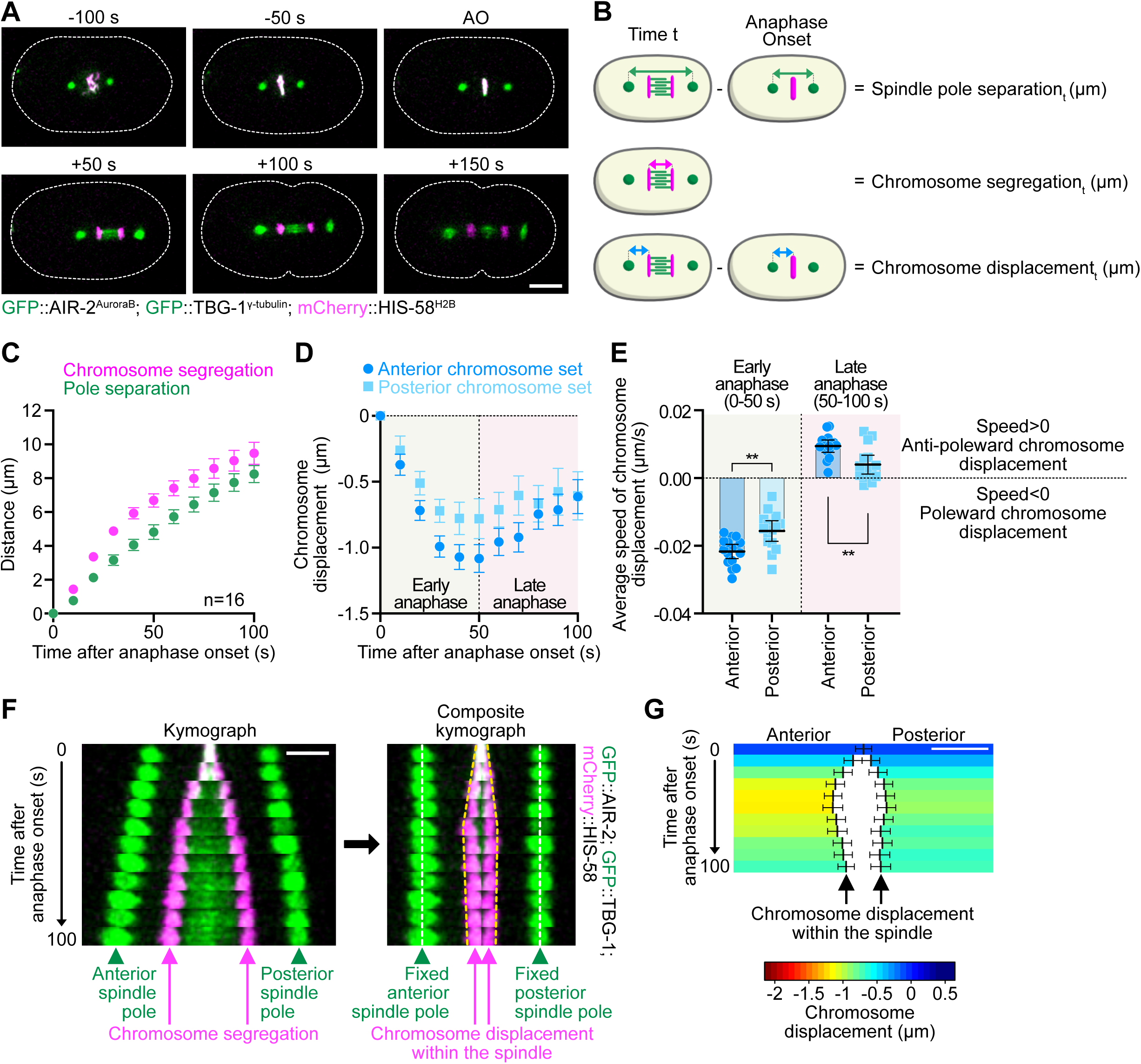
Chromosomes undergo a biphasic movement during anaphase. **(A)** Representative time-lapse images of a control zygote expressing mCherry-HIS-58^H2B^ (magenta), GFP::TBG-1^γ-tubulin^ (green) and GFP::AIR-2^AuroraB^ (green) during mitosis. Timings indicated above each image are from anaphase onset (AO). The dotted white line shows the outline of the zygote. Embryos are oriented with their antero-posterior axis aligned along the left-right direction. Scale bar, 10 µm. **(B)** Workflow for measuring spindle pole separation (green), chromosome segregation (magenta), and chromosome displacement (blue) overt time during anaphase. (P=Pole, C=Chromosome, AO=anaphase onset). **(C)** Quantification of the chromosome segregation (magenta) and spindle pole separation (green) over time from anaphase onset in control zygotes (n=16). **(D)** Quantification of the mean chromosome displacement over time from anaphase onset for the anterior (dark blue circles) and posterior (light blue squares) chromosome sets (n=16). **(E)** Quantification of average speed of chromosome displacement during early (0-50 seconds after anaphase onset) and late (50-100 seconds after anaphase onset) anaphase for the anterior (dark blue circles) and posterior (light blue squares) chromosome sets. Negative values (speed < 0) indicate a poleward chromosome displacement, positive values (speed > 0) indicate an anti-poleward chromosome displacement (n=16 zygotes). Mann-Whitney test on the mean speed of chromosome displacement at the anterior and posterior (**p<0.01). **(F)** Left: Kymograph aligned to the spindle center at 10-second intervals from a control zygote expressing mCherry-HIS-58^H2B^ (magenta), GFP::TBG-1^γ-tubulin^ (green), and GFP::AIR-2^AuroraB^ (green), recorded from 0 to 100 seconds post-anaphase onset. Right: Corresponding composite kymograph with the position of spindle poles fixed, as detailed in **Fig. S1 A**. White and yellow dotted lines indicate spindle poles and chromosome positions, respectively. Scale bar: 5 µm. **(G)** Color-coded graph representation of chromosome displacement for the anterior (left) and posterior (right) chromosome sets during anaphase. Scale bar, 2 µm. The color code is indicated below the graph. All error bars represent the 95% confidence interval.

We next measured the distance between each chromosome set and its respective spindle pole over time. We observed a biphasic pattern of chromosome displacement (**Fig. 1 D**). During the first 50 seconds after anaphase onset (early anaphase), we observed a decrease in the chromosome-to-pole distance, consistent with poleward chromosome movement and the existence of anaphase A. This was followed by a behavioral shift between 50 and 100 seconds after anaphase onset (late anaphase), where the chromosome sets reversed their movement relative to their respective spindle poles, moving anti-poleward (**Fig. 1 D**). Chromosome displacement was slower during this second anti-poleward phase than during the initial anaphase A-like phase (**Fig. 1 E**). Chromosome displacement was also evident in composite kymographs, where we artificially fixed the distance between the spindle poles over time (**Fig. 1 F**; and **Fig. S1 A**). To further emphasize the contribution of chromosomal displacement to overall chromosome segregation, independent of spindle pole separation, we generated color-coded graphs (**Fig. S1 B**). These graphs depicted chromosome displacement relative to their metaphase position, with a gradient from dark blue (anti-poleward movement) to dark red (strong poleward movement) (**Fig. 1 G**). The graphs also highlighted the asymmetric nature of chromosome displacement, which was more pronounced and faster for the anterior chromosome set compared to the posterior set (**Fig. 1, F and G**). Importantly, we observed similar results across three different *C. elegans* strains with varying genetic backgrounds (**Fig. S1, C and D**). Overall, our findings reveal a previously overlooked, reproducible, anaphase A-like mechanism of chromosome segregation during the first embryonic division in *C. elegans*. This chromosome movement follows a biphasic asymmetric pattern, transitioning from poleward displacement in early anaphase to anti-poleward movement in late anaphase.

#### Chromosome displacement is regulated by asymmetric cortical pulling forces

Next, we investigated the origin of the asymmetric behavior of the anterior and posterior chromosome sets during their displacement (**Fig. 1, D-G**). In the *C. elegans* zygote, the spatial segregation of PAR polarity proteins (PAR-3/PAR-6/PKC-3 in the anterior and PAR-1/PAR-2 in the posterior) drives the posterior enrichment of GPR-1/2^PINS^ and LIN-5^NuMA^, which act as cortical receptors for the microtubule minus-end-directed motor dynein (**Fig. 2 A**) (Delattre and Goehring, 2021; Lang and Munro, 2017; Rose and Gonczy, 2014). Cortical dynein, also enriched in the posterior, then generates asymmetric pulling forces on astral microtubules emanating from the two opposite spindle poles, ultimately leading to the posterior shift of the zygotic spindle and the first asymmetric division (Grill et al., 2001). We tested whether the asymmetric chromosome displacement observed in control zygotes was downstream of embryonic polarity. For this, we analyzed the behavior of spindle poles and chromosomes following RNAi-mediated depletion of polarity proteins (**Fig. 2**). In the absence of PAR-2, PAR-3 localizes uniformly across the cell cortex, resulting in uniformly low levels of cortical GPR-1/2^PINS^, LIN-5^NuMA^ and dynein (Colombo et al., 2003; Etemad-Moghadam et al., 1995). Conversely, in PAR-3-depleted zygotes, PAR-2 is uniformly localized to the cortex, which leads to a higher uniform distribution of cortical GPR-1/2^PINS^, LIN-5^NuMA^ and dynein (Boyd et al., 1996; Gotta et al., 2003; Park and Rose, 2008; Tsou et al., 2003). As expected, in both conditions, as well as upon RNAi-mediated depletion of GPR-1/2^PINS^, the zygotic spindle remained centrally positioned in the zygote and failed to migrate toward the posterior pole as observed in controls (**Fig. 2 B**). Interestingly, anaphase A chromosome poleward movement was also symmetric when cortical pulling forces were symmetrized (**Fig. 2, C and D**; and **Fig. S2 A**). Consistently, the difference in the speed of chromosome displacement between the anterior and posterior set of chromosomes observed in controls during both early and late anaphase, was entirely abolished in these symmetrized zygotes (**Fig. 2 E**). Interestingly, we also observed that in zygotes depleted of PAR-2 and GPR-1/2^PINS^, the biphasic pattern of chromosome displacement differed significantly from control zygotes. Instead of the typical transition from an initial poleward movement to a subsequent anti-poleward shift, chromosomes either remained static (*i.e*., showed no movement relative to the spindle poles) in the absence of PAR-2, or continued their poleward movement at a reduced speed in the absence of GPR-1/2^PINS^ (**Fig. 2 F**; and **Fig. S2 A**). In PAR-3-depleted embryos, spindle pole separation and the biphasic chromosome displacement profile were comparable to controls, consistent with the loss of the LET-99 lateral band in this condition, which prevents excessive cortical pulling forces (**Fig. 2, D and F-G**) (Tsou et al., 2002). Taken together, these results demonstrate that chromosome displacement in anaphase A is regulated downstream of polarity cues by the imbalance in cortical pulling forces between the anterior and posterior of the zygote during anaphase.

**Figure 2.**
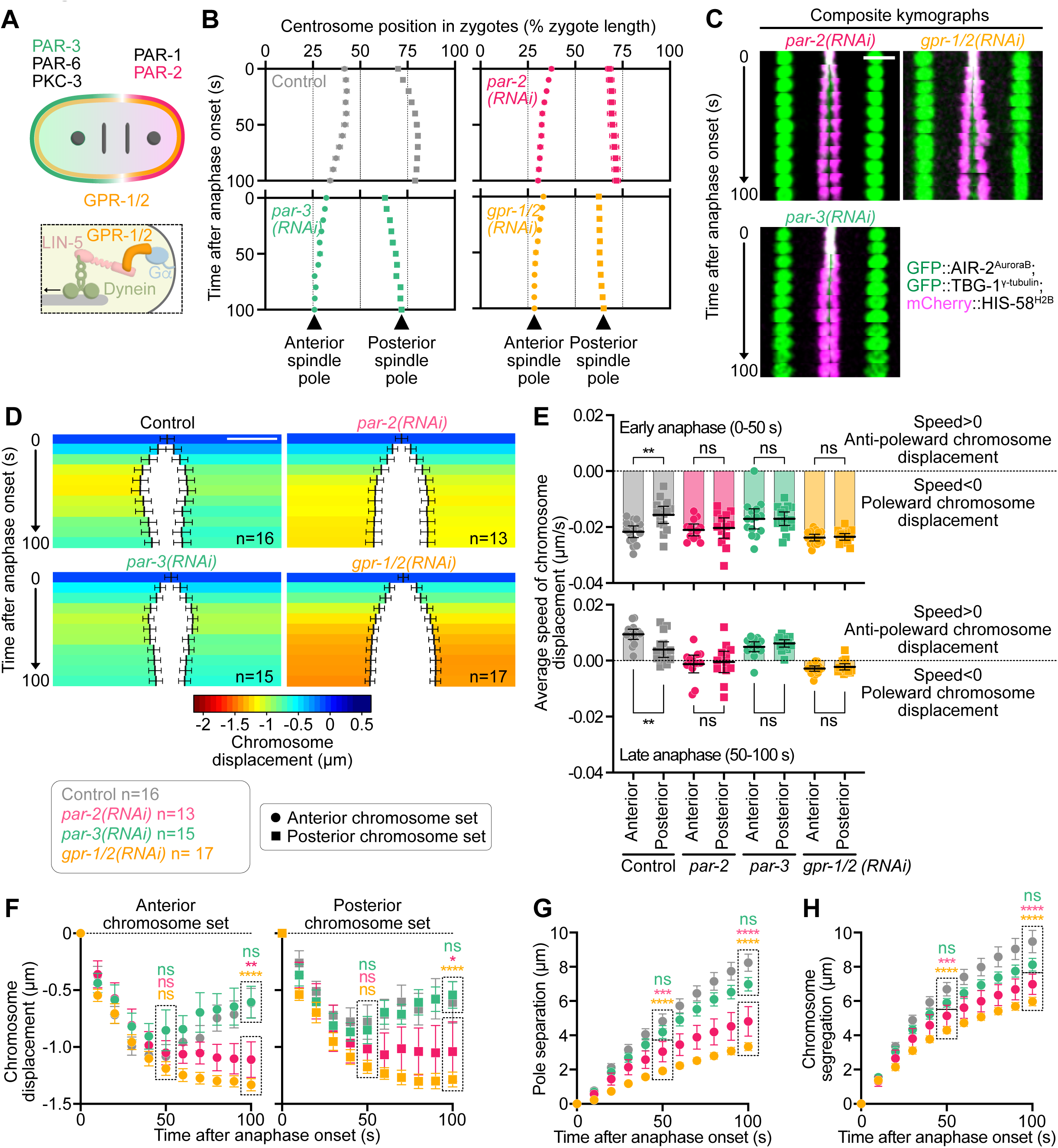
Chromosome displacement is regulated by asymmetric cortical pulling forces. **(A)** Top: Schematic representation of polarity determinant localization at the *C. elegans* zygote cortex. Bottom: Close-up illustrating the cortical dynein receptor at the *C. elegans* zygote cortex. **(B)** Quantification of the average relative position of spindle poles from the anterior (0%) to the posterior cortex (100%) of the zygote during anaphase in the indicated conditions (control n=16, *par-2(RNAi)* n=13, *par-3(RNAi)* n=15 and *gpr-1/2^PINS^(RNAi)* n=17 zygotes). Circles represent the anterior centrosome, squares represent the posterior centrosome. **(C)** Composite kymographs at 10-second intervals from zygotes expressing mCherry-HIS-11^H2B^ (magenta), GFP::TBG-1^γ-tubulin^ (green) and GFP::AIR-2^AuroraB^ (green), recorded from 0 to 100 seconds post-anaphase onset. Scale bar, 5 µm. **(D)** Color-coded graph representations of chromosome displacement for the anterior (left) and posterior (right) chromosome sets during anaphase. Scale bar, 2 µm. The color code is indicated below the graph. The sample size is indicated at the bottom right corner of each color-coded graph. **(E)** Quantification of average speed of chromosome displacement during early (0-50 seconds after anaphase onset) and late (50-100 seconds after anaphase onset) anaphase for the anterior (circles) and posterior (squares) chromosome sets. Negative values (speed<0) indicate a poleward chromosome displacement, positive values (speed>0) indicate an anti-poleward chromosome displacement. The sample size is the same as in (B). Mann-Whitney test on the mean speed of chromosome displacement at the anterior and posterior (ns: non-significant, **p<0.01). **(F)** Quantification of the mean chromosome displacement over time from anaphase onset for the anterior (top, circles) and posterior (bottom, squares) chromosome sets. The sample size is the same as in (B). One-way ANOVA Kruskal-Wallis with Dunn’s correction test (ns: non-significant, *p<0.05, **p<0.01, ****p<0.0001). **(G-H)** Quantification of average spindle pole separation (G) and chromosome segregation (H) over time from anaphase onset in indicated conditions. One-way ANOVA Kruskal-Wallis with Dunn’s correction test (ns: non-significant, ***p<0.001, ****p<0.0001). All error bars represent the 95% confidence interval.

### Anaphase A-like poleward chromosome displacement is negatively regulated by mechanical tension within the spindle

The lack of anti-poleward chromosome displacement in zygotes depleted of PAR-2 or GPR-1/2^PINS^ was associated with reduced spindle pole separation (**Fig. 2 G**) —a proxy for the cortical pulling forces acting on the spindle (Grill et al., 2001)—and by reduced chromosome segregation (**Fig. 2 H**). This finding implies a potential link between the mechanical tension within the spindle and chromosome displacement during anaphase. To further investigate this hypothesis, we disrupted the central spindle (or spindle midzone), a structure that typically withstands cortical pulling forces and maintains high mechanical tension within the spindle after anaphase onset (Grill et al., 2001; Raich et al., 1998; Verbrugghe, 2004). We previously demonstrated that the conserved antiparallel microtubule crosslinker SPD-1^PRC1^ is crucial for maintaining the mechanical integrity of the anaphase central spindle in *C. elegans* zygotes (**Fig. 3 A**) (Edwards et al., 2015; Hirsch et al., 2022; Maton et al., 2015). Accordingly, following RNAi-mediated depletion of SPD-1^PRC1^, we observed that spindle poles and chromosomes separated faster than in control zygotes (**Fig. 3, B and C**; **Fig. S2 B**; and **Video 2**). Consistent with our hypothesis, late anti-poleward chromosome displacement did not occur under this condition, with chromosomes instead continuing their poleward motion at a reduced speed during late anaphase (**Fig. 3, D and F**). Interestingly, the initial poleward chromosome displacement occurred faster in these zygotes compared to controls (**Fig. 3 G**). To confirm that the observed phenotypes were a consequence of reduced mechanical tension due to central spindle breakage, rather than to a specific function of SPD-1^PRC1^, we employed laser-mediated ablation to mechanically sever the central spindle. Importantly, this approach yielded identical results, with faster spindle pole separation correlating with both accelerated and sustained poleward chromosome displacement throughout anaphase (**Fig. 3, E-F**; and **Video 2**). Thus, reduction of cortical pulling forces (in PAR-3- or GPR-1/2^PINS^-depleted zygotes, **Fig. 2**) or the loss of central spindle-mediated resistance to these forces both lead to faster and sustained poleward chromosome displacement throughout anaphase (**Fig. 3**). Collectively, these findings demonstrate that poleward anaphase A-like chromosome displacement is restrained by mechanical tension within the spindle.

**Figure 3.**
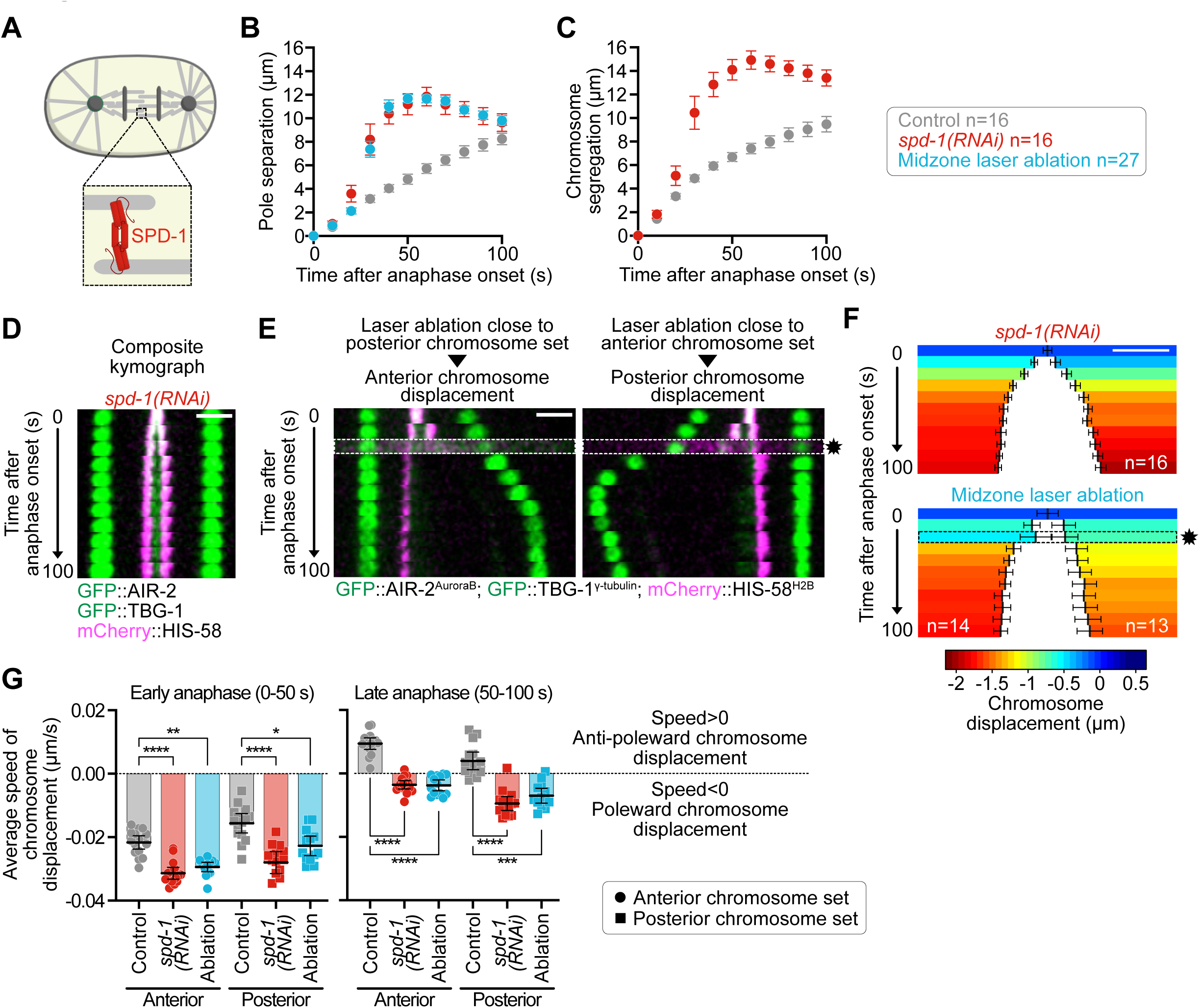
Anaphase A-like poleward chromosome displacement is regulated by tension within the spindle. **(A)** Schematic representation of the microtubule crosslinker SPD-1^PRC1^ at the anaphase central spindle in a *C. elegans* zygote. **(B-C)** Quantification of average spindle pole separation (B) and chromosome segregation (C) over time from anaphase onset in indicated conditions. (control n=16, *spd-1^PRC1^(RNAi)* n=16, midzone laser ablation n=27 zygotes). **(D)** Composite kymographs at 10-second intervals from a *spd-1^PRC1^(RNAi)* zygote expressing mCherry-HIS-58^H2B^ (magenta), GFP::TBG-1^γ-tubulin^ (green) and GFP::AIR-2^AuroraB^ (green), recorded from 0 to 100 seconds post-anaphase onset. Scale bar, 5 µm. **(E)** Kymographs aligned to the anterior (left, posterior ablation) or posterior (right, anterior ablation) spindle pole at 10-second intervals from midzone-ablated zygotes expressing mCherry-HIS-58^H2B^ (magenta), GFP::TBG-1^γ-tubulin^ (green) and GFP::AIR-2^AuroraB^ (green), recorded from 0 to 100 seconds post-anaphase onset. The midzone ablation timing (approximately 20 seconds after anaphase onset) is indicated by a black star and white dotted box. Scale bar, 5 µm. **(F)** Color-coded graph representations of chromosome displacement for the anterior (left) and posterior (right) chromosome sets during anaphase. Scale bar, 2 µm. The color code is indicated below the graph. The sample size is indicated at the bottom of each color-coded graph. **(G)** Quantification of average speed of chromosome displacement during early (0-50 seconds after anaphase onset) and late (50-100 seconds after anaphase onset) anaphase for the anterior (circles) and posterior (squares) chromosome sets. Negative values (speed<0) indicate a poleward chromosome displacement, positive values (speed>0) indicate an anti-poleward chromosome displacement. The sample size is the same as in (B). Mann-Whitney test on the mean speed of chromosome displacement at the anterior and posterior (*p<0.05, **p<0.01, ***p<0.001, ****p<0.0001). All error bars represent the 95% confidence interval.

### The depolymerizing kinesin KLP-7^MCAK^ is the primary driver of anaphase A-like poleward chromosome displacement

To identify the drivers of anaphase A-like chromosome displacement within the spindle, we conducted a targeted mini-screen in which mitotic microtubule-severing and kinesin-like proteins were co-depleted with GPR-1/2^PINS^ (**Fig. 4 A**; and **Fig. S2 C**). Depleting GPR-1/2^PINS^ induces persistent anaphase A-like movement throughout anaphase (**Fig. 2**) and minimizes spindle tension perturbations associated with depletion of candidate proteins, thereby facilitating clearer interpretation of results. We focused on proteins capable of depolymerizing, severing, or sliding microtubules, and present during mitosis in the *C. elegans* zygote. Our screen included the microtubule-severing proteins fidgetin FIGL-1^FIGN^ and spastin SPAS-1^SPAST^, the depolymerizing kinesin-13 KLP-7^MCAK^, the homotetrameric kinesin-5 BMK-1^Eg5^, the kinesin-6 family member ZEN-4^MKLP-1^, the kinesin-12 KLP-18, the minus-end-directed kinesin-14 KLP-15/16, and the chromokinesin-4 KLP-12 (Ali et al., 2000; Bishop et al., 2005; Matsushita-Ishiodori et al., 2007; Raich et al., 1998; Segbert et al., 2003; Srayko et al., 2005; Taguchi et al., 2022; Yakushiji et al., 2004). With the notable exceptions of KLP-7^MCAK^ and KLP-18, all co-depletion conditions phenocopied the single depletion of GPR-1/2^PINS^. This phenotype was characterized by faster and prolonged anaphase A-like poleward chromosome displacement compared to controls, indicating that these factors likely do not contribute to the regulation of chromosome displacement and anaphase A (**Fig. 4 A**; and **Fig. S2 C**). In contrast, the co-depletion of KLP-18 with GPR-1/2^PINS^ exacerbated this phenotype, resulting in an even greater anaphase A-like poleward displacement of chromosomes (**Fig. 4, A-F**). This suggests that KLP-18 acts as a negative regulator of anaphase A-like poleward chromosome displacement. Interestingly, co-depletion of KLP-7^MCAK^ produced the opposite effect, almost completely blocking anaphase A-like poleward chromosome displacement (**Fig. 4, A-D**) and reducing chromosome segregation, even though pole separation was slightly enhanced (**Fig. 4, E and F**). Importantly, we observed similar effects following single depletions of KLP-18 or KLP-7^MCAK^ in the presence of GPR-1/2^PINS^, although with higher variability likely due to the impact of depleting these proteins on cortical pulling forces and/or central spindle integrity when GPR-1/2^PINS^ are present (**Fig. S2, D-F**) (Han et al., 2015; Segbert et al., 2003; Srayko et al., 2005). To validate the role of KLP-7^MCAK^ in chromosome displacement, we used CRISPR/Cas9 to delete the entire coding sequence of klp-7^MCAK^, generating a *Δklp-7^MCAK^* mutant strain (**Fig. S2, G-I**). Consistent with prior findings, we confirmed that worms lacking KLP-7^MCAK^ are viable when maintained below 24°C (Gigant et al., 2017). Importantly, GPR-1/2^PINS^ depletion in the *Δklp-7^MCAK^* strain also blocked anaphase-A chromosome displacement. Together, our findings demonstrate that anaphase A-like poleward chromosome displacement requires the microtubule-depolymerizing kinesin-13 KLP-7^MCAK^ and is opposed by the kinesin-12 KLP-18.

**Figure 4.**
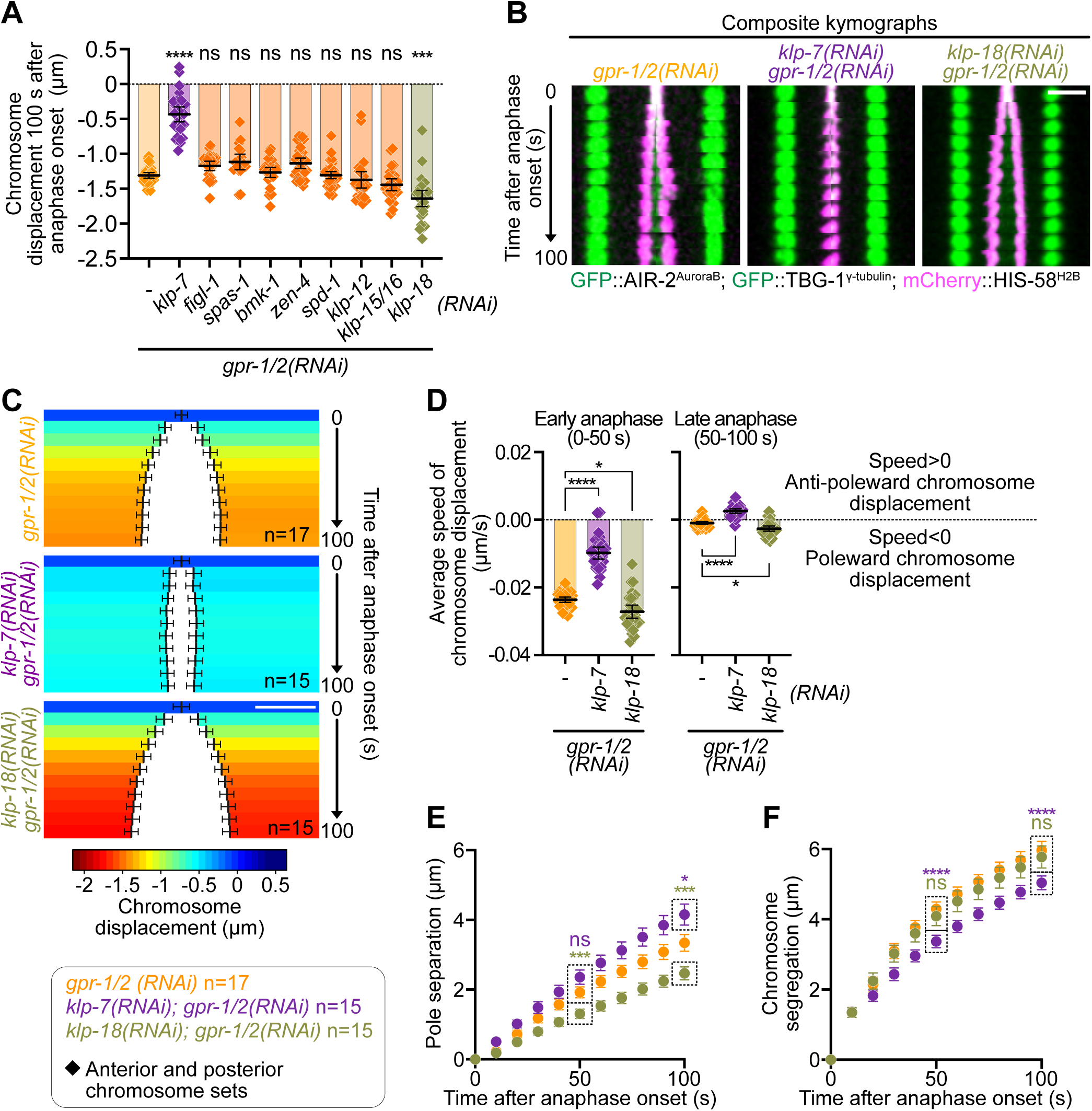
The depolymerizing kinesin KLP-7^MCAK^ drives anaphase A-like chromosome displacement. **(A)** Quantification of chromosome displacement 100 seconds after anaphase onset in indicated conditions (*gpr-1/2(RNAi)* n=17, klp-7(RNAi)*; gpr-1/2(RNAi)* n=15, *figl-1(RNAi); gpr-1/2(RNAi)* n=13, *spas-1(RNAi); gpr-1/2(RNAi)* n=10, *bmk-1(RNAi); gpr-1/2(RNAi)* n=14, *zen-4(RNAi); gpr-1/2(RNAi)* n=16, *spd-1(RNAi); gpr-1/2(RNAi)* n=21, *klp-12(RNAi); gpr-1/2(RNAi)* n=14, *klp-15/16(RNAi); gpr-1/2(RNAi)* n=14, *klp-18(RNAi); gpr-1/2(RNAi)* n=15 zygotes. ANOVA Kruskal-Wallis test comparing the single *gpr-1/2(RNAi)* to all double RNAi conditions (ns: non-significant, ***p<0.001, ****p<0.0001). **(B)** Composite kymographs at 10-second intervals from indicated conditions in zygotes expressing (A) mCherry-HIS-58^H2B^ (magenta), GFP::TBG-1^γ-tubulin^ (green) and GFP::AIR-2^AuroraB^ (green) recorded from 0 to 100 seconds post-anaphase onset. Scale bar, 5 µm. **(C)** Color-coded graph representations of chromosome displacement for the anterior (left) and posterior (right) chromosome sets during anaphase. The color code is indicated below the graph. The sample size is indicated at the bottom of each color-coded graph. Scale bar, 2 µm. **(D)** Quantification of average speed of chromosome displacement during early (0-50 seconds after anaphase onset) and late (50-100 seconds after anaphase onset) anaphase. Negative values (speed<0) indicate a poleward chromosome displacement, positive values (speed>0) indicate an anti-poleward chromosome displacement. Sample sizes are indicated below the graph and are the same as in (C). One-way ANOVA Kruskal-Wallis test (ns: non-significant, *p<0.05, ****p<0.0001). **(E-F)** Quantification of average spindle pole separation (E) and chromosome segregation (F) over time from anaphase onset in indicated conditions. One-way ANOVA Kruskal-Wallis with Dunn’s correction test (ns: non-significant, *p<0.05, ***p<0.001, ****p<0.0001). All error bars represent the 95% confidence interval.

### Anaphase A-like poleward chromosome displacement increases during early embryonic development

Successive cell cleavage divisions during early embryonic development produce progressively smaller cells, accompanied by a proportional decrease in spindle size (**Fig. S3 A**) (Hara and Kimura, 2009; Lacroix and Dumont, 2022; Lacroix et al., 2018; Reber and Goehring, 2015; Rieckhoff et al., 2020). To determine whether the chromosome displacement observed within the spindle during the zygotic division persists in later embryonic divisions, we analyzed *C. elegans* embryos from the 1-cell to the 64-cell stage (**Fig. 5 A**; **Fig. S3, B and C**; and **Video 3**). As expected, chromosome segregation and spindle pole separation decreased proportionally with cell size (Hara and Kimura, 2009; Lacroix et al., 2018). However, the contribution of anaphase-A poleward chromosome displacement within the spindle to overall chromosome segregation increased progressively across successive divisions, reaching up to 49.9±2.9% at the 64-cell stage (**Fig. 5, B-D**). Furthermore, the extent of poleward chromosome displacement 100 seconds after anaphase onset inversely correlated with spindle pole separation, a readout of cortically generated tension on the spindle (**Fig. 5 E**). In conclusion, our results show that as blastomeres decrease in size during early embryonic divisions, the contribution of anaphase A–like poleward chromosome displacement to overall segregation becomes increasingly pronounced. We propose that this progressive enhancement of anaphase A likely reflects a reduction in spindle tension in smaller blastomeres.

**Figure 5.**
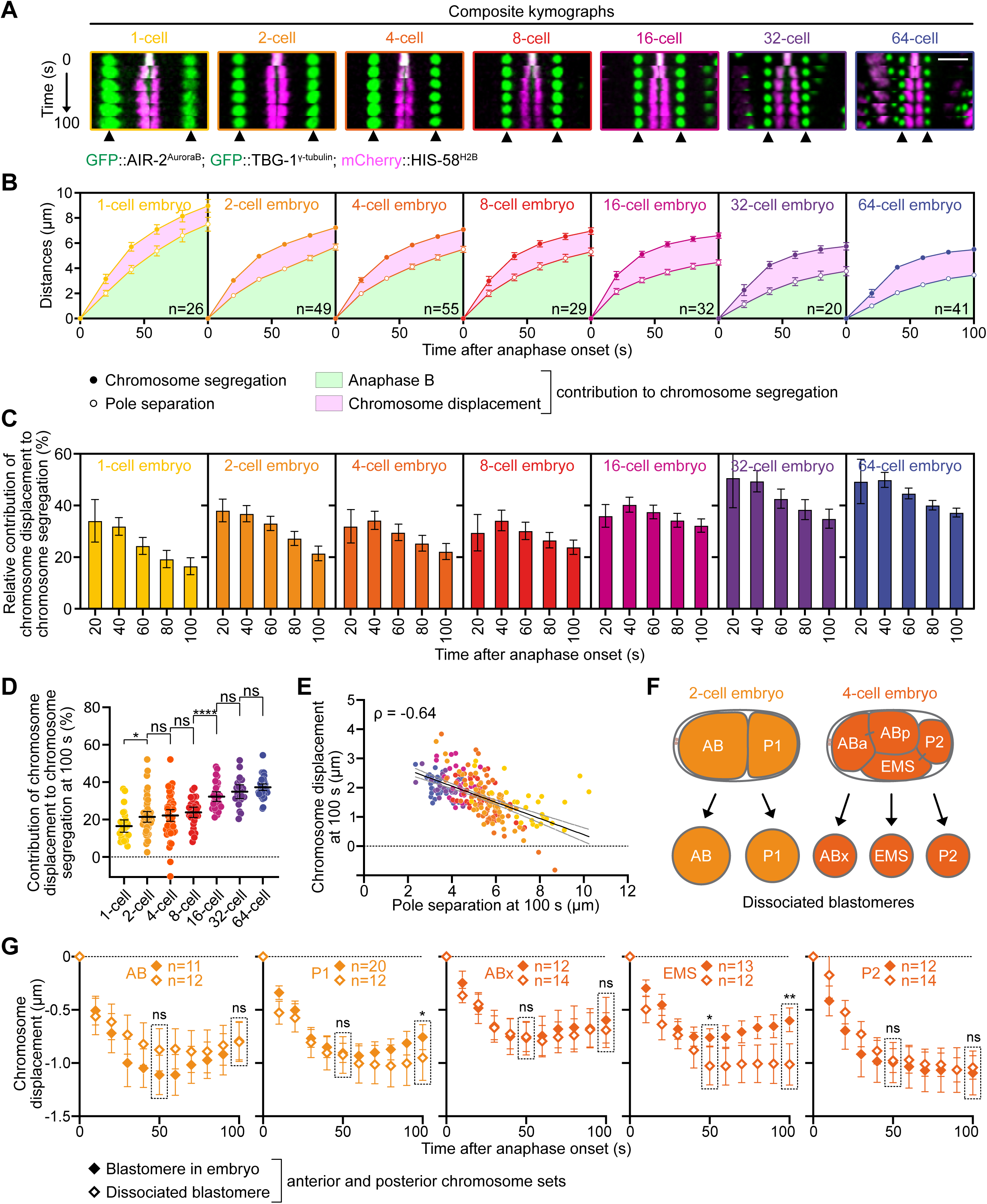
Anaphase A-like poleward chromosome displacement increases during early embryonic development. **(A)** Composite kymographs at 20-second intervals from indicated developmental stages in embryos expressing mCherry-HIS-58^H2B^ (magenta), GFP::TBG-1^γ-tubulin^ (green) and GFP::AIR-2^AuroraB^ (green), recorded from 0 to 100 seconds post-anaphase onset. Arrowheads at the bottom indicate the aligned spindle poles. Scale bar, 5µm. **(B)** Quantification of average chromosome segregation (filled circles) and spindle pole separation (open circles) over time from anaphase onset in indicated developmental stages (1-cell n=26, 2-cell n=49, 4-cell n=55, 8-cell n=29, 16-cell n=32, 32-cell n=20, and 64-cell n=41 embryos). Green areas represent chromosome segregation due to spindle pole separation (Anaphase B), pink areas represent chromosome segregation due to chromosome displacement. **(C)** Quantification of the relative contribution of chromosome displacement to overall chromosome segregation over time during anaphase at each developmental stage. Sample sizes are the same as in (B). **(D)** Quantification of the average relative contribution of chromosome displacement to overall chromosome segregation 100 seconds after anaphase onset at each developmental stage. Mann-Whitney test (ns: non-significant, *p<0.05, ****p<0.0001). The color code is the same as in (C). Sample sizes are the same as in (B). **(E)** Plot of chromosome displacement over spindle pole separation 100 seconds after anaphase onset. The color code is the same as in (C). Sample sizes are the same as in (B). Black and grey dotted lines represent the linear regression and 95% confidence interval respectively. Spearman correlation coefficient is indicated at the top left corner (ρ=-0.64, p<0.0001). **(F)** Schematic representation of the blastomere organization in the 2-cell (left) and 4-cell embryo (right), and after blastomere isolation. **(G)** Quantification of the mean chromosome displacement over time from anaphase onset for blastomeres in intact embryos (filled diamonds) or after blastomere isolation (open diamonds). Sample sizes are indicated at the top of each graph. Mann-Whitney test (ns: non-significant, *p<0.05, **p<0.01). All error bars represent the 95% confidence interval.

Finally, we examined whether chromosome displacement within the spindle in multicellular embryos is regulated cell-autonomously, through intracellular mechanisms controlling mechanical tension within the spindle, or non-autonomously, via cell-cell communication or contact-induced mechanical tension (di Pietro et al., 2016; Lisica et al., 2022; van Leen et al., 2020). To test this, we performed blastomere dissociation on 2- and 4-cell embryos (**Fig. 5 F**; and **Video 4**) and compared the variation in chromosome-to-pole distance in dissociated blastomeres with that of their corresponding control non-dissociated blastomeres (**Fig. S3, D-E**; and **Fig. 5 G**). Interestingly, chromosome displacement was comparable between control and dissociated blastomeres in all cases, with the notable exception of the 4-cell stage EMS blastomere (Sulston et al., 1983). In dissociated EMS blastomeres (**Fig. 5 G**, open diamonds), chromosome movement toward the spindle poles was significantly prolonged and lacked the subsequent anti-poleward displacement observed in control 4-cell stage embryos (**Fig. 5 G**, filled diamonds). EMS cells become polarized through physical contact with P2 neighbors (Goldstein, 1992; Goldstein and Hird, 1996). This EMS-P2 contact activates the Wnt and MES-1/SRC-1 pathways in EMS, modulating cortical pulling forces that are essential for proper EMS spindle orientation and asymmetric division (Rose and Gonczy, 2014). Thus, this result further supports the idea that cell polarity and asymmetric cortical pulling forces regulate chromosome displacement within the spindle during anaphase. Overall, we conclude that chromosome displacement is primarily governed by intrinsic cell-autonomous mechanisms. However, in the 4-cell stage EMS blastomere, extrinsic, cell-cell contact-dependent mechanisms play a crucial role in controlling the extent of anaphase A-like chromosome displacement and facilitating the transition to the anti-poleward phase of chromosome movement.

### Conclusion

Overall, our results uncover the existence of a transient KLP-7^MCAK^-mediated anaphase A, which plays an increasingly significant role in chromosome segregation during early *C. elegans* development. We propose that this anaphase A is inhibited by tension within the spindle resulting from astral microtubule cortical pulling forces resisted by the anaphase central spindle (Grill et al., 2001). These cortical pulling forces normally promote anaphase B. Thus, our results highlight an unexpected mechanical coordination between anaphase A and B in early embryos.

During anaphase, KLP-18 is enriched in the central spindle region, where it is thought to mediate microtubule crosslinking and sliding (Segbert et al., 2003; Tanenbaum and Medema, 2010). Thus, it remains unclear whether KLP-18 directly inhibits anaphase A, or whether its depletion destabilizes the central spindle, thereby reducing spindle tension and indirectly promoting anaphase A. Our observation that depleting KLP-18 in absence of GPR-1/2^PINS^ led to increased poleward chromosome displacement coupled with decreased spindle pole separation during anaphase (**Fig. 4, E and F**) is consistent with a recent report in human HeLa cells, suggesting that anaphase A can potentially restrict spindle pole separation rather than promote chromosome segregation (Chen et al., 2025). However, our results following KLP-7^MCAK^ perturbations (**Fig. 4, E and F**), which show a strong decrease in chromosome segregation with minimal effect on spindle pole separation, demonstrate that, unlike in HeLa cells, the primary effect of anaphase A in *C. elegans* embryos is to directly drive chromosome segregation.

Like its vertebrate counterpart MCAK, KLP-7^MCAK^ localizes to both kinetochores and spindle poles in *C. elegans* embryos (Encalada et al., 2005; Gigant et al., 2017; Han et al., 2015; Oegema et al., 2001; Schlaitz et al., 2007), potentially promoting kinetochore microtubule shortening from both sites. However, how spindle tension modulates KLP-7^MCAK^ activity during anaphase remains unclear. In vertebrates, MCAK activity and localization are regulated in a tension-dependent manner by centrosomal Aurora A and kinetochore-localized Aurora B providing a potential mechanism for the differential regulation of various microtubule populations (Andrews et al., 2004; Lan et al., 2004; Ohi et al., 2004; Zhang et al., 2008). Similarly, AIR-1^Aurora^ ^A^ and/or AIR-2^Aurora^ ^B^-dependent phospho-regulation of KLP-7^MCAK^ activity could respond to anaphase spindle tension and underlie the fine-tuning of anaphase chromosome displacement in *C. elegans* early embryos (Han et al., 2015). Loss of KLP-7^MCAK^ may also increase other spindle microtubule populations, such as central spindle microtubules, so the effects of depletion (**Fig. 4, E and F**) may reflect these changes, rather than anaphase A alone.

The reversal of chromosome movement from poleward to anti-poleward during anaphase that we observed here appears to be a rare and largely overlooked phenomenon, with only a few prior reports in other systems (Skibbens et al., 1993; Wurzenberger et al., 2012). While the underlying mechanisms remain to be elucidated, this switch could arise from a shift in microtubule dynamics—from depolymerization to polymerization—potentially governed by a balance between kinase and phosphatase activities directed toward microtubule dynamics regulators (Wurzenberger et al., 2012). In *C. elegans* zygotes, kinetochore microtubules are formed mainly of multiple overlapping short segments rather than continuous fibers extending from kinetochore to centrosome (Redemann et al., 2017), raising the possibility that microtubule sliding driven by kinesin-like motor proteins also contributes to this unusual reversal.

We further propose that chromosome movement in each half-spindle are differentially regulated by cell polarity cues, which are known to asymmetrically control astral microtubule–mediated cortical pulling forces (Labbe et al., 2003). Interestingly, asymmetric anaphase A has also been observed in maize meiosis, although the underlying molecular mechanisms in that system remain unclear (Nannas et al., 2016). Our observation that the contribution of anaphase A progressively increases across successive divisions in a largely cell-autonomous manner further suggests that spindle tension decreases throughout successive early divisions. This progressive reduction in spindle tension is likely driven by the gradual shortening of astral microtubules, and thus by the corresponding decrease in length-dependent pulling forces, observed over successive divisions (Grill et al., 2003; Kimura and Onami, 2005; Kozlowski et al., 2007; Lacroix et al., 2018). We propose that the increased contribution of anaphase A serves as a compensatory mechanism to maintain robust chromosome segregation when pulling forces weaken. This interpretation aligns with our model of coordinated anaphase A and B contributions and provides a developmental rationale for the shift toward enhanced anaphase A. Altogether, our results highlight a previously uncharacterized coordination between anaphase A and B, which participates in sister chromatid physical separation.

## MATERIAL AND METHODS

### *C. elegans* strains and Maintenance

The list of *C. elegans* strains employed in this study can be found in **Table S1**. Strains were maintained at 23°C on Nematode Growth Medium plates (NGM, 51 mM NaCl, 2.5 g Bacto Peptone, 17 g Bacto Agar, 12 µM Cholesterol, 1 mM CaCl_2_, 1 mM MgSO_4_, 25 µM KH_2_PO_4_ and 5 µM Nystatin) seeded with *E. coli* OP50 bacteria (Brenner, 1974). Transgenic lines were generated through crossing pre-existing strains or engineered by CRISPR/Cas9 mutagenesis (details below) or Mos1-mediated Single Copy Insertion (MosSCI) (Frokjaer-Jensen, 2015). All worms analyzed were hermaphrodites. Wormbase.org was used as a resource throughout this work (Sternberg et al., 2024).

### RNA Interference

Double stranded RNAs (dsRNAs) were synthesized from DNA templates that were PCR-amplified using primers flanked with T3 or T7 phage promoter sequences (listed in **Table S2**). PCR products were purified (PCR purification kit, Qiagen). The purified DNA templates were then used for *in vitro* T3 and T7 RNA transcription reactions (Megascript, Invitrogen, #AM1334 for T7 and #AM1338 for T3) for 5 hours at 37°C. Following transcription, RNAs were purified (MEGAclear kit, Invitrogen, #AM1908), denatured for 10 minutes at 68°C, and annealed 30 minutes at 37°C. Aliquots of 2 µL were flash frozen in liquid nitrogen and stored at -80°C. L4 larvae were injected at the specified concentration and incubated at 20°C for 44-48 hours before imaging.

### Generation of the *klp-7^MCAK^* deletion by CRISPR/Cas9

A *klp-7^MCAK^* deletion *C. elegans* strain was generated using CRISPR/Cas9. The Cas9-NLS purified protein (MacroLab Facility, University of California Berkeley), along with tracrRNAs and crRNAs (Integrated DNA Technologies, Inc.) were mixed for 15 minutes at 37°C before injection in the JDU570 strain. The crRNAs were designed using the Alt-R HDR Design Tool (IDT website). The injection mix contained *klp-7^MCAK^*specific crRNAs, namely *crJD13* (5’-acttttcatcgggatcgaat-3’) and *crJD14* (5’-aagtgggtagcatatcgtcg-3’) designed to cut before the ATG start and after the TGA stop codon, respectively. A *dpy-10* crRNA (5‘-gctaccataggcaccacgag-3’) and a repair template generating the *dpy-10(cn64)* mutation (cacttgaacttcaatacggcaagatgagaatgactggaaaccgtaccgcatgcggtgcctatggtagcggagcttcacatggcttcagaccaacagcctat) by homologous recombination-based repair served as a co-injection marker (Arribere et al., 2014). The *klp-7^MCAK^* deletion was generated through Non-Homologous End-Joining (NHEJ) without a repair template. 92 hours after injection, roller (*dpy-10(cn64)*/*dpy-10(+)* and dumpy (*dpy-10(cn64)*/*dpy-10(cn64)*) worms were isolated on NGM plates. The presence of the *klp-7^MCAK^* deletion was confirmed through single worm PCR. For this, individual adult worms were lyzed in 2.5 µL Worm Lysis Buffer (0.5 mg/mL Proteinase K (Promega, V3021) in 1X ThermoPol Buffer (New England BioLabs, B9004S)), for 1 hour at 65°C. Proteinase K was then heat-inactivated for 15 minutes at 95°C, and PCR was subsequently performed using Taq DNA Polymerase with ThermoPol Buffer (NEB, M0267) with *oJD943* (5’-ttcccactccatcgttgattgg-3’) and *oJD1097* (5’-tagaatgtttttgttaaatgcgatacg-3’) primers to amplify the *klp-7^MCAK^* deleted allele, or *oJD807* (5’-acattttcagggcgagacaa-3’) and oJD808 (5’-tgtggttgatggagaattgtg-3’) to amplify *klp-7^MCAK^* wild-type allele. Sequencing revealed that the *klp-7^MCAK^* deletion extends from 13 nucleotides prior to the ATG start codon to 24 nucleotides following the TGA stop codon.

### Live Imaging and kymographs

Fluorescently labeled adult worms were dissected in 6 µL Meiosis Medium (5 mg/mL inulin, 25 mM HEPES, 60% Leibovitz’s L-15 median and 20% fetal bovine serum). The extracted embryos were mounted between two coverslips and sealed with a Vaseline petroleum jelly gasket (Laband et al., 2018).

For imaging isolated blastomeres, young gravid hermaphrodites were dissected in ddH_2_O and the eggshell was subsequently removed by an alkaline hypochlorite treatment. Embryos were then transferred to Shelton’s growth medium (0.52x *Drosophila* Schneider’s Medium, 0.288 mg/mL inulin, 2.88 mg/mL Polyvinylpolypyrrolidone, 0.0059x BME vitamins, 0.0059x chemically defined lipid concentrate, 0.59x Penn-Strep, and 0.35x FBS). The vitelline envelope was removed, and the blastomeres were dissociated through repeated aspiration and ejection using a 30 µm diameter glass needle (World Precision Instruments). Blastomeres were then mounted in 20 µL of Shelton’s growth medium before imaging (Davies et al., 2018). Imaging of embryos was conducted at 23°C using a Nikon Ti-E inverted microscope equipped with a Yokogawa CSU-X1 spinning-disk confocal head, with a Nikon APO λS 60x/NA1.4 oil objective, and a CoolSNAP HQ2 CCD camera (Photometrics Scientific) with 2×2 binning. The temperature was maintained at 22-23°C using a thermostatic chamber enclosing the microscope. Dissociated blastomeres and their respective 2- and 4-cell stage control embryos were imaged using a Nikon Ti inverted microscope equipped with a Yokogawa CSU-10 spinning disk confocal head, with Borealis (Spectral Applied Research) and an Orca-R2 charge-coupled camera (Hamamatsu Photonics). In all cases, acquisition parameters were controlled with the Metamorph 7 software (Molecular Devices, RRID:SCR_002368).

For zygotes (**Fig. 1–5**), 4 Z-sections spaced 2 µm apart were captured every 10 seconds. For multi-cell embryos (**Fig. 5**), imaging parameters varied based on cell stage and spindle orientation, including: 4 Z-sections at 2 µm every 10 seconds, 9 Z-sections at 2 µm every 10 seconds, 40 Z-sections at 1 µm every 20 seconds, and 70 Z-sections at 0.5 µm every 20 seconds. Despite these variations, all parameters produced consistent results. Therefore, all acquired movies were included in the subsequent analysis and kymographs, which were generated at 20 second intervals.

Embryos displayed in **Fig. 5 A**, were imaged using the following acquisition settings: 1-cell P0, 2-cell AB, 2-cell P1, 4-cell EMS, 4-cell P2, 8-cell and 16-cell embryos were acquired with 4 Z-sections spaced 2 µm apart at 10 second intervals, 4-cell Abd, 32-cell and 64-cell embryos were imaged with 70 Z-sections spaced 0.5 µm apart at 20 second intervals. Composite kymographs were generated at 20 second intervals. For embryos analyzed in **Fig. 5 G** and illustrated in **Fig. S3, D-E**, images were acquired with 9 Z-plans spaced 2 µm apart at 10 second intervals.

All kymographs were generated using Fiji (RRID:SCR_002285) (Schindelin et al., 2012) and Affinity Designer 2 (Affinity, RRID:SCR_016952) softwares. Raw images were pre-filtered with a 0.5-pixel mean filter in Fiji. The final images are maximal Z-projections.

Composite kymographs were specifically crafted to emphasize the contribution of chromosome displacement to chromosome segregation, excluding the contribution of spindle pole separation. This process involved producing individual kymographs aligned on the anterior or posterior spindle pole for each embryo. The two kymographs were vertically cropped at the midpoint of chromosomes at anaphase onset and displayed side-by-side **(Fig. S1 A)**.

### Anaphase central spindle laser ablation

Central spindle laser ablation was conducted using the iLas Pulse system (Roper Scientific) with a switched 355 nm UV-pulsed laser with a repetition rate of 6 kHz, integrated into our spinning-disk microscope (Laband et al., 2017). Beginning 20 seconds after anaphase onset, a series of 100–130 UV pulses were systematically delivered along a 1-pixel-wide line oriented perpendicular to the central spindle. To avoid photobleaching of both sets of segregating chromosomes, the ablation line was positioned proximal to the anterior or posterior sets of segregating chromosomes 20 seconds after anaphase onset.

### Chromosome and spindle pole tracking and quantitative analysis

Chromosome and spindle pole tracking analysis were conducted using Imaris software (Oxford Instruments, RRID:SCR_007370). The Surface function in Imaris was used for semi-automatic tracking of chromosomes and spindle poles during the 150 second interval following anaphase onset. Spatial coordinates of chromosomes and spindle pole centers were extracted over time, allowing calculation of Euclidean distances: between the spindle poles (PP), between the two segregating chromosome sets (CC), and between each chromosome set and its respective spindle pole (CP) in the anterior or posterior spindle halves. The distances were calculated using the formula:

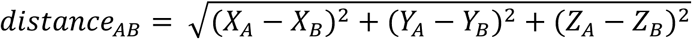

The spindle pole separation (ΔPP) and chromosome displacement (ΔCP) throughout anaphase were determined by subtracting the measured distances at a given time (t) to the corresponding distance at anaphase onset (AO):

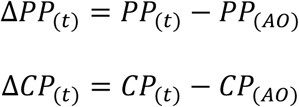

The chromosome segregation (CC) is the Chromosome-to-Chromosome distance at a given timepoint (t).

The velocity of chromosome-to-pole movement during early anaphase (0 to 50 seconds after anaphase onset) and late anaphase (50 to 100 seconds after anaphase onset) was calculated using the formulas:

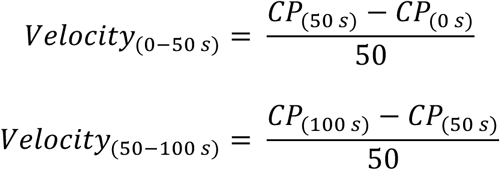

The direction of movement was determined by the velocity sign: a negative sign indicated poleward chromosome movement, while a positive sign represented anti-poleward movement.

The overall contribution of Chromosome-to-Pole distance variation (ΔCP) at a given timepoint was the difference between the spindle pole separation (ΔPP) and the chromosome segregation (CC). This contribution, expressed as a percentage of chromosome segregation at that timepoint, was calculated using this equation:

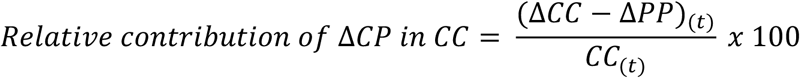

### Generation of color-coded graphs

Color-coded graphs illustrating Chromosome-to-Pole distance variation (ΔCP) relative to anaphase onset in the anterior and posterior regions of the mitotic spindle were generated using a custom Python script (RRID:SCR_008394; https://github.com/SergeDmi/spindle_colors). In the graphs, vertical black lines represented the segregating chromosome masses, while the white segments in between depicted chromosome segregation driven by Chromosome-to-Pole distance variation. Graphs were cropped to account for spindle size variations across different experimental conditions, ensuring a consistent and standardized visual representation.

### Figure preparation, graphs and statistical analyses

Figures and illustrations were crafted using Affinity Designer 2 software, while all graphs (excluding color-coded graphs) and statistical analysis were conducted with Prism 10 (GraphPad, RRID:SCR_002798). Each statistical test employed is outlined in the respective figure legend.

## SUMMARY OF SUPPLEMENTAL MATERIAL

3 supplementary figures: Fig. S1 describes the method we used to generate the color-coded graphs and composite kymographs. Fig. S2 shows the effect of depleting polarity proteins, microtubule crosslinkers and motors on chromosome displacement. Fig. S3 displays the dynamics of spindle pole elongation and chromosome segregation from the one to the 64-cell stage in *C. elegans* embryos.

4 supplementary videos: Video 1 is from Fig. 1A-C. Video 2 is from Fig. 3. Video 3 is from Fig. 5A-E. Video 4 is from Fig. 5F, G.

2 supplementary tables: Table S1, *C. elegans* strains used in this study. Table S2, oligonucleotides used in this study.

## DATA AVAILABILITY STATEMENT

All data supporting the findings of this study are available within the paper and its Supplementary Information.

## ACKNOWLEDGEMENTS

We thank all members of the Dumont lab for support and advice. We are grateful to Patricia Moussounda, Clarisse Picard, and Téo Bitaille for providing technical support. We thank Isabelle Bécam for her input on the manuscript. This work was supported by CNRS and University Paris Cité, by NIH R01GM117407 and R01GM130764 (J.C. Canman), and by grant from the European Research Council ERC-CoG ChromoSOMe 819179 (J. Dumont).

## SUPPLEMENTARY FIGURE LEGENDS

Figure S1. **Workflow for generating composite kymographs and color-coded graphs. (A)** Top left: kymograph, aligned on the anterior spindle pole, at 10-second intervals from a zygote expressing mCherry-HIS-58^H2B^ (magenta), GFP::TBG-1^γ-tubulin^ (green) and GFP::AIR-2^AuroraB^ (green), recorded from 0 to 100 seconds post-anaphase onset. Top right: Corresponding kymograph aligned on the posterior spindle pole. Middle: Cropped kymographs along the anterior (left) and posterior (right) set of segregating chromosomes. Bottom: Composite kymograph generated by joining the cropped kymographs to allow visualizing chromosome displacement. Scale bar, 5 µm. **(B)** Workflow for generating the color-coded graphs. For each timepoint (from 0 to 100 seconds after anaphase onset), the chromosome displacement was color-coded from dark blue (0.5 µm anti-poleward displacement) to red (2 µm poleward displacement) and mounted vertically as a kymograph, with the anterior pole on the left and the posterior pole on the right. The width of the uncropped color-coded graph indicates the spindle length at anaphase onset. For easy comparison, all color-coded graphs were arbitrarily cropped to the same width. Chromosomes are depicted by vertical black lines. Chromosome segregation driven by chromosome displacement is visualized as the white space between black lines. **(C-D)** Composite kymographs (C) and their corresponding color-coded graphs (D) for zygotes expressing TBG-1^γ-tubulin^::mScarlet and mCherry::HIS-11^H2B^ (left), TBG-1^γ-tubulin^::mCherry and mCherry::HIS-58^H2B^ (middle) and TBG-1^γ-tubulin^::GFP and GFP::HIS-11^H2B^ (right). The sample size is indicated at the bottom of each color-coded graph. Scale bar, 2 µm. All error bars represent the 95% confidence interval.

Figure S2. **Targeted mini-screen to identify proteins involved in regulating chromosome displacement. (A)** Quantification of the mean chromosome displacement over time from anaphase onset for the anterior (circles) and posterior (squares) chromosome sets (control n=16, *par-2(RNAi)* n=13, *par-3(RNAi)* n=15, *gpr-1/2^PINS^(RNAi)* n=17 embryos). **(B)** Quantification of the average relative position of spindle poles from the anterior (0%) to the posterior cortex (100%) of the zygote during anaphase in the indicated conditions (*spd-1^PRC1^(RNAi)* n=16, midzone laser ablated n=27 zygotes). Circles represent the anterior centrosome, squares represent the posterior centrosome. **(C-D)** Color-coded graph representations of chromosome displacement for the anterior (left) and posterior (right) chromosome sets during anaphase in indicated conditions. Scale bar, 2 µm. The color code is indicated under (D). The sample size is indicated at the bottom right corner of each color-coded graph. **(E-F)** Quantification of average spindle pole separation (E) and chromosome segregation (F) over time from anaphase onset in indicated conditions. One-way ANOVA Kruskal-Wallis with Dunn’s correction test (ns: non-significant, **p<0.01, ***p<0.001. **(G)** Composite kymographs at 10-seconds interval from indicated conditions in zygotes expressing TBG-1^γ-tubulin^::mScarlet and mCherry-HIS-11^H2B^ (grey), recorded from 0 to 100 seconds post-anaphase onset. Scale bar, 5 µm. **(H)** Color-coded graph representations of chromosome displacement for the anterior (left) and posterior (right) chromosome sets during anaphase. The color code is the same as in (C). The sample size is indicated at the bottom of (E). Scale bar, 2 µm. **(I)** Quantification of average speed of chromosome displacement during early (0-50 seconds after anaphase onset) and late (50-100 seconds after anaphase onset) anaphase. Negative values (speed<0) indicate a poleward chromosome displacement, positive values (speed>0) indicate an anti-poleward chromosome displacement. Sample sizes are the same as in (F). Mann-Whitney test (ns: non-significant, ****p<0.0001). All error bars represent the 95% confidence interval.

Figure S3. **Chromosome displacement during early embryogenesis in control or isolated blastomeres. (A)** Quantification of spindle length at anaphase onset plotted at each cleavage stage (1- to 64-cell stage). **(B,C)** Quantification of chromosome segregation (B) and spindle pole elongation (C) over time during anaphase at each cleavage stage (1- to 64-cell stage). Sample sizes and the color code for (A-C) are indicated on the right. **(D-E)** Composite kymographs at 20-second intervals from indicated developmental stages in embryos expressing mCherry-HIS-58^H2B^ (magenta), GFP::TBG-1^γ-tubulin^ (green) and GFP::AIR-2^AuroraB^ (green) for control (D) or dissociated (E) blastomeres, recorded from 0 to 100 seconds post-anaphase onset. Arrowheads at the bottom indicate the aligned spindle poles. Scale bar, 5µm. All error bars represent the 95% confidence interval.

## VIDEO LEGENDS

**Video 1 is related to Fig. 1, A, B and C. Four-dimensional tracking of chromosomes and spindle poles during mitosis in *C. elegans* zygote.** (top) 3D projection of a time-lapse movie of a control zygote expressing mCherry-HIS-58^H2B^ (magenta), GFP::TBG-1^γ-tubulin^ (green) and GFP::AIR-2^AuroraB^ (green) during mitosis and (bottom) the corresponding tracking of chromosomes (cyan) and spindle poles (yellow). Images were acquired every 10 second with 9 Z-plans separated by 2 µm for each timepoint. Timings indicated are related to anaphase onset. Scale bar 10 µm. Playback speed, 7 frames/second.

**Video 2 is related to Fig. 3. Anaphase A chromosome displacement after SPD-1^PRC1^ depletion or midzone laser ablation.** Time-lapse imaging of *C. elegans* zygotes expressing mCherry-HIS-58^H2B^ (magenta), GFP::TBG-1^γ-tubulin^ (green) and GFP::AIR-2^AuroraB^ (green) during mitosis in control condition (top left), after RNAi-mediated depletion of SPD-1^PRC1^ (top right), or after midzone laser ablation performed approximately 20 seconds after anaphase onset, near the posterior set of chromosomes (bottom left) or near the anterior set of chromosomes (bottom right). Movies were acquired every 10 second and are maximum projections of 4 Z-plans separated by 2 µm for each timepoint. Timings indicated are related to anaphase onset. Scale bar 10 µm. Playback speed, 7 frames/second.

**Video 3 is related to Fig. 5, A-E. 4D tracking of chromosomes and spindle poles at the 32-cell stage.** 3D projection of a time-lapse movie of a control 32-cell embryo expressing mCherry-HIS-58^H2B^ (magenta), GFP::TBG-1^γ-tubulin^ (green) and GFP::AIR-2^AuroraB^ (green) during mitosis and (bottom) the corresponding tracking of chromosomes (cyan) and spindle poles (yellow). Images were acquired every 20 seconds with 72 Z-plans separated by 0.5 µm for each timepoint. Timings indicated are related to anaphase onset. Scale bar 10 µm. Playback speed, 7 frames/second.

**Video 4 is related to Fig. 5, F and G. Chromosome segregation in 2-cell stage dissociated blastomeres.** Time-lapse imaging of a 2-cell *C. elegans* embryo (top) or dissociated blastomeres AB and P1 (bottom) expressing mCherry-HIS-58^H2B^ (magenta), GFP::TBG-1^γ-tubulin^ (green) and GFP::AIR-2^AuroraB^ (green) during mitosis Movies were acquired every 10 second and are maximum projections of 4 Z-plans separated by 2 µm for each timepoint. Scale bar 10 µm. Playback speed, 7 frames/second.

## Notes

### Competing Interest Statement

The authors have declared no competing interest.

