## Supplementary figures and tables for "Mechanical coordination between anaphase A and B drives asymmetric chromosome segregation"

### Figure S1

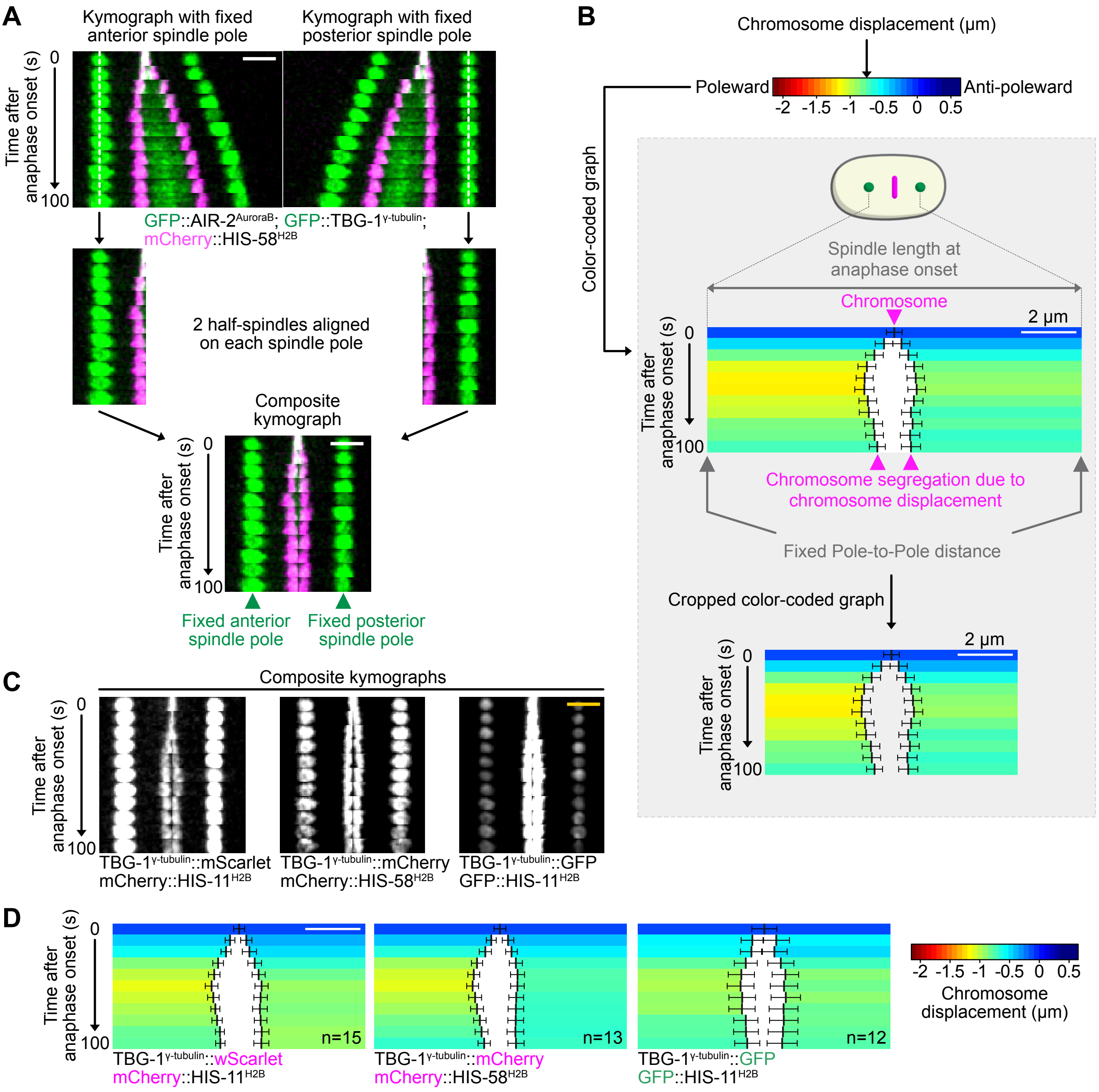

A
Kymograph with fixed anterior spindle pole
Kymograph with fixed posterior spindle pole
Time after anaphase onset (s)
0
100
GFP::AIR-2AuroraB; GFP::TBG-1γ-tubulin;
mCherry::HIS-58H2B
2 half-spindles aligned on each spindle pole
Composite kymograph
Fixed anterior spindle pole
Fixed posterior spindle pole
B
Chromosome displacement (μm)
Poleward
Anti-poleward
-2 -1.5 -1 -0.5 0 0.5
Color-coded graph
Spindle length at anaphase onset
Chromosome
2 μm
Chromosome segregation due to chromosome displacement
Fixed Pole-to-Pole distance
Cropped color-coded graph
C
Composite kymographs
TBG-1γ-tubulin::mScarlet
mCherry::HIS-11H2B
TBG-1γ-tubulin::mCherry
mCherry::HIS-58H2B
TBG-1γ-tubulin::GFP
GFP::HIS-11H2B
D
n=15
n=13
n=12
TBG-1γ-tubulin::wScarlet
mCherry::HIS-11H2B
TBG-1γ-tubulin::mCherry
mCherry::HIS-58H2B
TBG-1γ-tubulin::GFP
GFP::HIS-11H2B
Chromosome displacement (μm)

### Figure S2

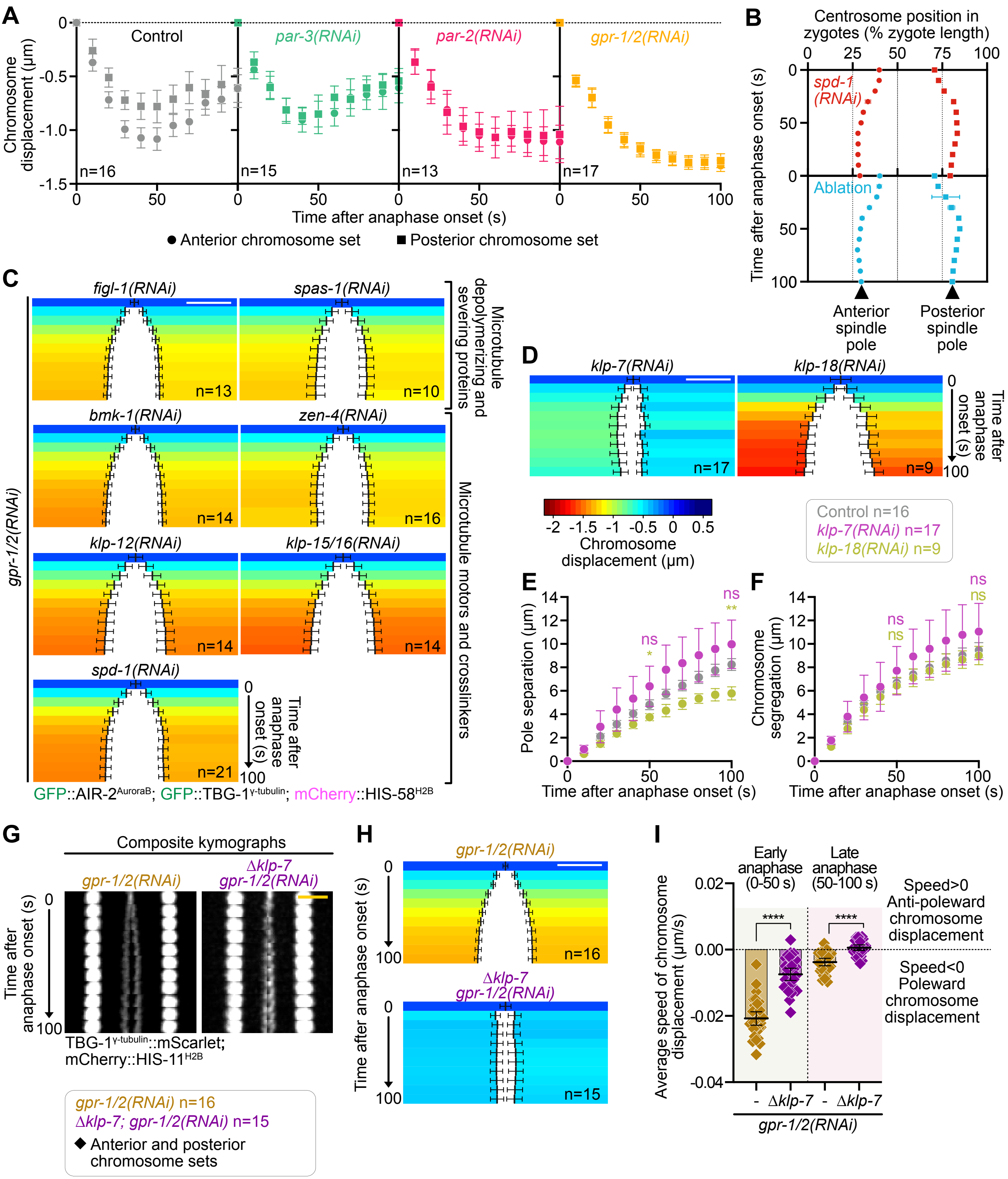

A
Chromosome displacement (µm)
Control
par-3(RNAi)
par-2(RNAi)
gpr-1/2(RNAi)
n=16
n=15
n=13
n=17
Time after anaphase onset (s)
Anterior chromosome set
Posterior chromosome set
B
Centrosome position in zygotes (% zygote length)
Time after anaphase onset (s)
spd-1 (RNAi)
Ablation
Anterior spindle pole
Posterior spindle pole
C
gpr-1/2(RNAi)
figl-1(RNAi)
spas-1(RNAi)
n=13
n=10
Microtubule depolymerizing and severing proteins
bmk-1(RNAi)
zen-4(RNAi)
n=14
n=16
klp-12(RNAi)
klp-15/16(RNAi)
n=14
n=14
Microtubule motors and crosslinkers
spd-1(RNAi)
n=21
Time after anaphase onset (s)
GFP::AIR-2AuroraB; GFP::TBG-1γ-tubulin; mCherry::HIS-58H2B
D
klp-7(RNAi)
klp-18(RNAi)
n=17
n=9
Time after anaphase onset (s)
Chromosome displacement (µm)
Control n=16
klp-7(RNAi) n=17
klp-18(RNAi) n=9
E
Pole separation (µm)
Time after anaphase onset (s)
ns
*
**
F
Chromosome segregation (µm)
Time after anaphase onset (s)
ns
G
Composite kymographs
gpr-1/2(RNAi)
Δklp-7 gpr-1/2(RNAi)
Time after anaphase onset (s)
TBG-1γ-tubulin::mScarlet; mCherry::HIS-11H2B
gpr-1/2(RNAi) n=16
Δklp-7; gpr-1/2(RNAi) n=15
Anterior and posterior chromosome sets
H
gpr-1/2(RNAi)
n=16
Δklp-7 gpr-1/2(RNAi)
n=15
Time after anaphase onset (s)
I
Early anaphase (0-50 s)
Late anaphase (50-100 s)
Average speed of chromosome displacement (µm/s)
****
Speed>0 Anti-poleward chromosome displacement
Speed<0 Poleward chromosome displacement
- Δklp-7 - Δklp-7
gpr-1/2(RNAi)

### Figure S3

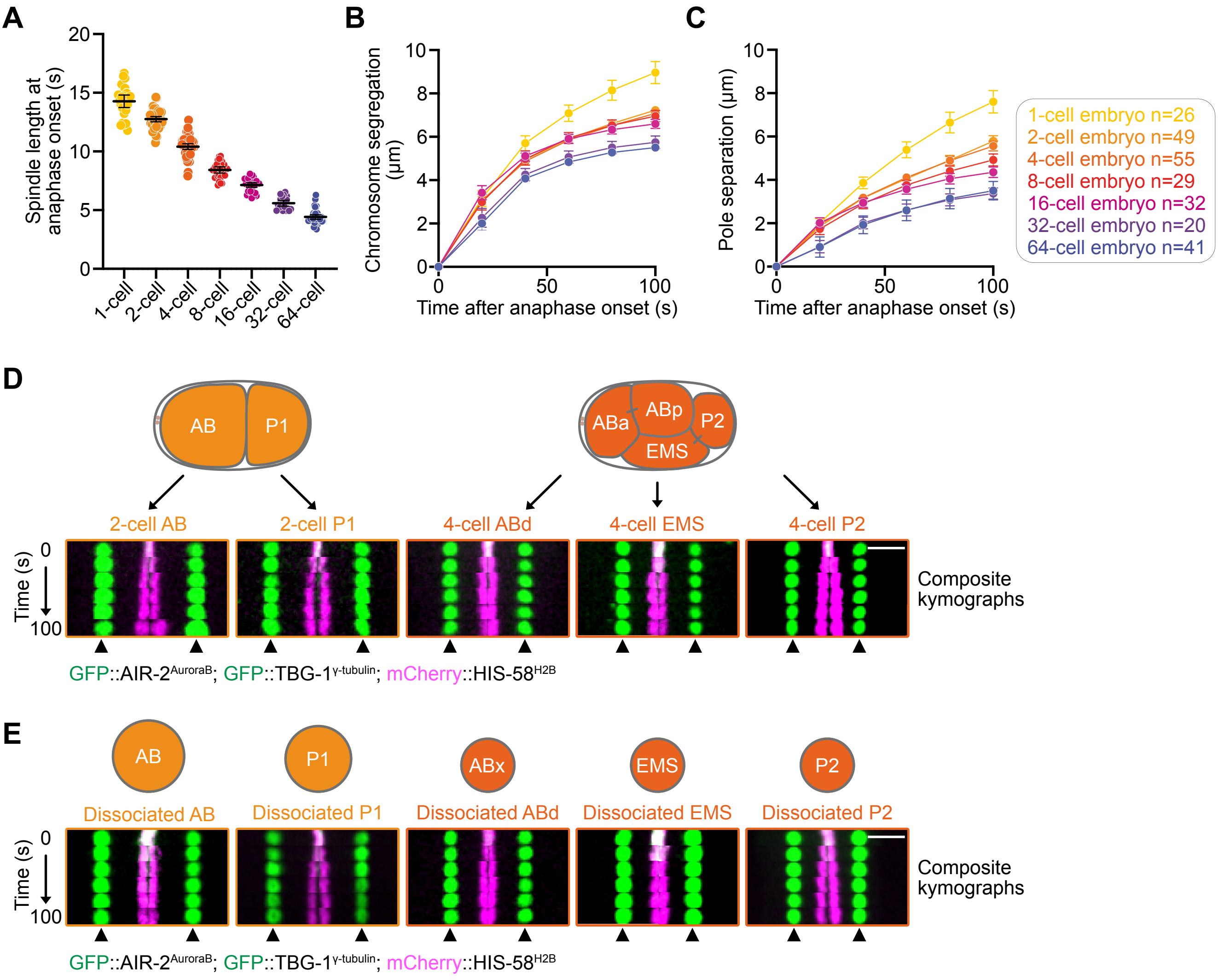

A
Spindle length at anaphase onset (s)
20
15
10
5
0
1-cell
2-cell
4-cell
8-cell
16-cell
32-cell
64-cell
B
Chromosome segregation (μm)
10
8
6
4
2
0
0
50
100
Time after anaphase onset (s)
C
Pole separation (μm)
Time after anaphase onset (s)
1-cell embryo n=26
2-cell embryo n=49
4-cell embryo n=55
8-cell embryo n=29
16-cell embryo n=32
32-cell embryo n=20
64-cell embryo n=41
D
AB
P1
ABa
ABp
P2
EMS
2-cell AB
2-cell P1
4-cell ABd
4-cell EMS
4-cell P2
Time (s)
0
100
Composite kymographs
GFP::AIR-2AuroraB; GFP::TBG-1γ-tubulin; mCherry::HIS-58H2B
E
AB
P1
ABx
EMS
P2
Dissociated AB
Dissociated P1
Dissociated ABd
Dissociated EMS
Dissociated P2
Time (s)
0
100
Composite kymographs
GFP::AIR-2AuroraB; GFP::TBG-1γ-tubulin; mCherry::HIS-58H2B

#### Supplementary Table 1: Strains used in this study

| Strain | Genotype | Source |
|---|---|---|
| JCC56 | unc-119(ed3)III?; ltIs37[pAA64; Ppie-1/mCherry::his-58; unc-119 (+)]IV, ltIs14[pASM05; Ppie-1/GFP-TEV-STag::air-2; unc-119 (+)]IV; ddIs6[Ppie-1/GFP:: tbg-1; unc-119(+)]V | Maton et al., 2015 |
| JDU570 | ijmSi125[pJD768; Pmex-5/tbg-1::mScarlet; mCherry::his-11; cb-unc-119(+)]II | This study |
| JDU585 | ijmSi125[pJD768; Pmex-5/tbg-1::mScarlet; mCherry::his-11; cb-unc-119(+)]II; klp-7(ijm13)III | This study |
| JDU711 | ltSi569[oxTi185; pOD1110/pSW008; CEOP3608 tbg-1::mCherry; cb-unc-119(+)]I; unc-119(ed3)III?; ltIs37[pAA64; pie-1/mCherry::his-58; unc-119 (+)]IV | This study |
| OD1702 | unc-119(ed3)III; ltSi560 [pPLG014; Pmex-5/GFP::his-11::tbb-2_3'UTR, tbg-1::gfp::tbb-2_3'UTR; cb-unc-119(+)]V | Kim et al., 2015 |

#### Table S2: Oligonucleotides used for dsRNA synthesis

| Gene | Oligonucleotide 1 (5' → 3') | Oligonucleotide 2 (5' → 3') | Template | [C] (mg/ml) |
|---|---|---|---|---|
| *spd-1* (Y34D9A.4) | TAATACGACTCACTATAGGctcttcccagtaaaggcgttcg | AATTAACCCTCACTAAAGGtttagccacgggctccatcttcg | gDNA | 1.28 |
| *gpr-1/2* (F22B7.13 / C38C10.4) | TAATACGACTCACTATAGGagcatgtgattccacacgtc | AATTAACCCTCACTAAAGGtctggcagcagacagttcag | gDNA | 1.74 |
| *par-2* (F58B6.3) | TAATACGACTCACTATAGGccggctccagagtgtcc | AATTAACCCTCACTAAAGGgccgtcgcccactgtcg | gDNA | 1.32 |
| *par-3* (F54E7.3) | TAATACGACTCACTATAGGagacttctagagatcaatgg | AATTAACCCTCACTAAAGGttgatgtgctgtggatcagc | gDNA | 0.91 |
| *figl-1* (F32D1.1) | TAATACGACTCACTATAGGaatggcagtacaacaatctcc | AATTAACCCTCACTAAAGGctcaatgtgaccagtggaatg | cDNA | 1.15 |
| *spas-1* (C24B5.2) | TAATACGACTCACTATAGGttagcaaccgaaacttcgagag | AATTAACCCTCACTAAAGGgcaagtcttccgatgcatcag | cDNA | 1.52 |
| *klp-7* (K11D9.1) | TAATACGACTCACTATAGGaaaaaggtgtggggcaagtt | AATTAACCCTCACTAAAGGgacacgggtgttcgagacta | cDNA | 1.30 |
| *bmk-1* (F23D12.8) | TAATACGACTCACTATAGGagctcaactgatgaccctac | AATTAACCCTCACTAAAGGgccatttcgcgaattcgatc | gDNA | 2.02 |
| *zen-4* (M03D4.1) | TAATACGACTCACTATAGGattggagctgttggatgagc | AATTAACCCTCACTAAAGGaattggttatggctccgaga | cDNA | 1.50 |
| *klp-12* (T01G1.1) | TAATACGACTCACTATAGGacactgaaactgaacgagatcg | AATTAACCCTCACTAAAGGttctacaatttccaccattac | gDNA | 1.14 |
| *klp-15/16* (M01E11.6 / C41G7.2) | TAATACGACTCACTATAGGtgcttgtcgtcctccgtctcgtttgc | AATTAACCCTCACTAAAGGtggattcagcgaagagaaatgcagc | gDNA | 1.31 |
| *klp-18* (C06G3.2) | TAATACGACTCACTATAGGgttgatgacctccgtgtcct | AATTAACCCTCACTAAAGGtgacgagaaacagaagtttgaca | gDNA | 1.19 |
